# Deep learning predicts tuberculosis drug resistance status from genome sequencing data

**DOI:** 10.1101/275628

**Authors:** Michael L. Chen, Akshith Doddi, Jimmy Royer, Luca Freschi, Marco Schito, Matthew Ezewudo, Isaac S. Kohane, Andrew Beam, Maha Farhat

**Affiliations:** Department of Biomedical Informatics, Harvard Medical School, Boston, MA; University of Virginia School of Medicine, Charlottesville, VA; Analysis Group Inc.; Critical Path Institute, 1730 E River Rd., Tucson, AZ; Division of Pulmonary & Critical Care, Massachusetts General Hospital, Boston, MA

## Abstract

**Background:** The diagnosis of multidrug resistant and extensively drug resistant tuberculosis is a global health priority. Whole genome sequencing of clinical *Mycobacterium* tuberculosis isolates promises to circumvent the long wait times and limited scope of conventional phenotypic antimicrobial susceptibility, but gaps remain for predicting phenotype accurately from genotypic data.

**Methods and Findings:** Using targeted or whole genome sequencing and conventional drug resistance phenotyping data from 3,601 *Mycobacterium tuberculosis* strains, 1,228 of which were multidrug resistant, we investigated the use of machine learning to predict phenotypic drug resistance to 10 anti-tuberculosis drugs. The final model, a multitask wide and deep neural network (MD-WDNN), achieved improved high predictive performance: the average AUCs were 0.979 for first-line drugs and 0.936 for second-line drugs during repeated cross-validation. On an independent validation set, the MD-WDNN showed average AUCs, sensitivities, and specificities, respectively, of 0.937, 87.9%, and 92.7% for first-line drugs and 0.891, 82.0% and 90.1% for second-line drugs. In addition to being able to learn from samples that have only been partially phenotyped, our proposed multidrug architecture shares information across different anti-tuberculosis drugs and genes to provide a more accurate phenotypic prediction. We use *t*-distributed Stochastic Neighbor Embedding (*t*-SNE) visualization and feature importance analyses to examine inter-drug similarities.

**Conclusions:** Machine learning is capable of accurately predicting resistant status using genomic information and holds promise in bringing sequencing technologies closer to the bedside.

## Introduction

Tuberculosis (TB) is among the top 10 causes of mortality worldwide with an estimated 10.4 million new incidents of TB in 2015 [1]. The growing use of antibiotics in healthcare has led to increased prevalence of drug resistant bacterial strains [2], and the World Health Organization (WHO) estimates that 4.1% of new *Mycobacterium tuberculosis* (MTB) clinical isolates are multidrug-resistant (MDR) (*i.e*. resistant to rifampicin [RIF] and isoniazid [INH]). Furthermore, approximately 9.5% of MDR cases are extensively drug-resistant (XDR) (*i.e*. resistant to one second-line injectable drug, such as amikacin [AMK], kanamycin [KAN], or capreomycin [CAP], and one fluoroquinolone, such as moxifloxacin [MOXI], or ofloxacin [OFLX]) [1]. The WHO estimates that 48% of MDR-TB and 72% of XDR-TB patients have unfavorable treatment outcomes, citing the lack of MDR-TB detection and treatment as a global health crisis [1].

Diagnosing drug resistance remains a barrier to providing appropriate TB treatment. Due to insufficient resources for building diagnostic laboratories, fewer than half of the countries with a high MDR-TB burden have modern diagnostic capabilities [3]. Even in the best equipped laboratories, conventional culture and culture based antimicrobial susceptibility testing constitutes a considerable biohazard and requires weeks to months before results are reported due to the slow *in vitro* growth of *Mycobacterium tuberculosis* [1]. Molecular diagnostics are now an increasingly common alternative to conventional cultures. The WHO has endorsed three such molecular tests: the GeneXpert MTB/RIF, a rapid RT-PCR based diagnostic test assay that detects RIF resistance, the Hain line probe assay (LPA) that tests for both RIF and INH resistance, and the Hain MDRTB*sl*, an LPA that tests for resistance to second-line injectable drugs and fluoroquinolones [1]. The LPAs approved by the WHO have seen moderate sensitivities, ranging from 63.7% to 94.4% for second-line injectable drugs and fluoroquinolones [4–6]. However, current diagnostic approaches face challenges. First, these methods have limited sensitivity because they rely on a few genetic loci, ranging between 1-6 loci per test [6,7]. Second, they do not detect most rare gene variants of the targeted loci, especially insertions, deletions, and variants in promoter regions [8]. Third, current molecular tests only detect resistance to five anti-tuberculosis drugs, notably missing several key first line agents, including pyrazinamide and ethambutol, and over 5 additional agents currently used for treatment. Fourth, they do not account for variables such as genetic background and gene-gene interactions despite good evidence for their contribution to resistance for several drugs including rifampicin, ethambutol and fluoroquinolones from allelic exchange experiments [9–11]. The limited scope of these tests suggests the need for a comprehensive antimicrobial susceptibility test.

An alternative to targeted mutation detection methods is whole genome sequencing, which captures both common and rare mutations involved in drug resistance. Past studies utilizing whole genome sequencing have shown a wide range of performance, with sensitivities for first-line drugs ranging from 54% to 98% [8,12,13]. Second-line injectable drugs and fluoroquinolones had lower sensitivities, most of which were between 30% and 96% [8,12,13]. We hypothesize that the limited predictive performance of anti-tuberculosis drugs outside of first-line drugs could be improved using a large dataset enriched for resistance to second-line drugs and a more complex model.

Deep learning models have become a powerful tool for many classification tasks. Modern deep neural networks have achieved state-of-the-art performance in image recognition [14], speech recognition [15], and natural language processing [16]. Researchers in medicine have begun to translate these approaches for use in personalized clinical care. Deep ‘convolutional’ neural networks have been used to in identifying diabetic retinopathy [17] and classifying skin cancers [18]. Deep learning applications in computational biology and bioinformatics have also been successful, such as in predicting RNA-binding protein sites [19], inferring target gene expression from landmark genes [20], and identifying biomarkers for predicting human chronological age [21]. The flexibility of deep learning architectures has allowed for a range of successful applications in clinical tasks, biomedicine, molecular genomics, and other fields.

We demonstrate here an improved predictive tool to evaluate drug resistance for 10 anti-tuberculosis drugs using a novel multidrug wide and deep neural network (MD-WDNN) framework [22]. In contrast to previously reported approaches that focus on predicting resistance a single drug at a time, our multidrug framework that predicts the full resistance profile simultaneously allows a drug to share resistance pathway information from the phenotypes of other drugs and incorporates prior knowledge that drug resistance can be caused by both direct genotype-phenotype relationships as well as epistatic effects [9–11]. We evaluate the deep learning architectural features to understand the relative influence of genomic markers, provide insights into the biological basis for our model, and gain a deeper understanding of the relationships amongst the 10 anti-tuberculosis drugs.

## Materials and Methods

### Experimental Design

MTB targeted sequence and antibiotic resistance data from a sample enriched in first and second-line antibiotic resistance [8] was pooled with public whole genome sequence and resistance data for training of the prediction model. Model validation was performed on an independent set of public whole genome sequences for which phenotypic resistance data was available. The validation dataset was a convenience dataset not preselected based on antibiotic resistance or strain lineage and diversity distribution. We evaluated MTB isolate diversity through hierarchical clustering and using lineage-defining mutations in the drug resistance loci, as assessed by Walker *et al*. [12]. In order to predict drug resistance for each isolate, we built a unified multidrug wide and deep neural network (MD-WDNN) to predict phenotypic status for all drugs simultaneously. For baselines, we compared the MD-WDNN to a random forest model and a logistic regression model with an L2 penalty (ridge regression). To understand how each component of the MD-WDNN contributed to its predictive performance, we performed an ablation study by comparing it to several variants in which a certain aspect was changed. These comparisons include a single-task WDNN where each drug was learned using a separate model (SD-WDNN), a MD-WDNN trained on only common mutations (e.g. no derived features), a multilayer perceptron without the ‘wide’ component (deep MLP), and a single-drug WDNN trained on mutations known to be resistance-determining for each drug [8] to evaluate the impact of training on the full genome sequence (kSD-WDNN). All comparisons were performed using 10-fold cross validation using the training data, repeated 5 times for a total for 50 evaluations of each model. We visualized the multidrug WDNN’s final phenotypic representation in 2-dimensional *t*-SNE plots and evaluated the importance of genetic variants to resistance through permutation testing.

### Data Description

#### Sequence data

The training dataset consisted of 1,379 MTB isolates that underwent sequencing using molecular inversion probes that targeted 28 preselected antibiotic resistance genes and promoter regions, with 100 bases flanking both ends of each region [8]. This sequence data was pooled with 2,222 additional MTB whole genome sequences curated by the ReSeqTB knowledgebase, which maintains a public data sharing platform (www.reseqtb.org) curating genotypic and phenotypic data of WHO-endorsed *in vitro* diagnostic assays for MTB [23]. The validation dataset of 792 MTB isolates was obtained by pooling additional data from ReSeqTB, without overlap with the training set, and other MTB whole genome sequences and phenotype data curated manually from the following references [24–27].

#### Antibiotic resistance phenotype data

All isolates included underwent culture based antibiotic susceptibility testing to two or more drugs at WHO approved critical concentrations and met other quality control criteria as detailed in [8]. The pooled phenotype data included resistance status for eleven drugs: first-line drugs (rifampicin, isoniazid, pyrazinamide, and ethambutol); streptomycin; second-line injectable drugs (capreomycin, amikacin, and kanamycin); and fluoroquinolones (ciprofloxacin, moxifloxacin, and ofloxacin). Phenotypic data was classified as resistant, susceptible, or not available.

### Variant calling

We used a custom bioinformatics pipeline to clean and filter the raw sequencing reads. We aligned filtered reads to the reference MTB isolate H37Rv using Stampy 1.0.23 [28] and variants were called by Platypus 0.5.2 [29] using default parameters. Genome coverage was assessed using SAMtools 0.1.18 [30] and read mapping taxonomy was assessed using Kraken [31]. Strains with a coverage of less than 95% at 10x or more in the regions of interest (Supplementary Table S6) or that had a mapping percentage of less than 90% to *Mycobacterium tuberculosis* complex were excluded. Further, regions of the remaining genome not covered by 10 regions or more in at least 95% of the isolates were filtered out from the analysis. In the remaining regions, variants were further filtered if they had a quality of <15, purity of <0.4, or did not meet the PASS filter designation by Platypus.

### Building the predictor set of features

Because 1,379 of the 3,601 of the MTB isolates in the training set underwent targeted sequencing only, we restricted the resistance predictors to variants in the regions targeted in these isolates (Supplementary Table S6). Since the *eis* and *rpsA* genes and promoters were recently determined to be associated with kanamycin and pyrazinamide resistance respectively [32,33], we added mutations in the *eis* and *rpsA* regions into our set of predictors. For those isolates with missing genotype data, we used a status of 0.5 for the missing mutations.

The features used for prediction consisted of two groups. In the first group, each mutation was considered a predictor and its status was binary (either present or absent). The second group, we created ‘derived’ categories by grouping the rarer mutations (present in <30 isolates) by gene locus (coding, intergenic and putative promoter regions). For each coding region, we split the variants by type into three groups: single nucleotide substitution (SNP), frameshift insertion/deletion, or non-frameshift insertion/deletion. For each non-coding region, we split the variants by type into two groups: insertions/deletion or single nucleotide substitution). We used individual and derived predictors found in at least 30 MTB isolates to make our final set of predictors used to train all models except kSD-WDNN and WDNN (Common Mutations).

### Evaluation of MTB isolate diversity

We identified lineage-defining variants as assessed in a 2015 study by Walker *et al*. [12]. The genetic-lineage similarity between each pair of isolates was computed as the Euclidean distance between the two corresponding lineage-defining mutation vectors. We applied Ward’s method of hierarchical clustering on the resultant distance matrix [34] to group the isolates and displayed the isolate-isolate Euclidean distance matrix based on the lineage-defining variants in a heat map. We used *hclust* in the R stats 3.4.2 package to perform hierarchical clustering. Each group was mapped back to the recognized MTB lineage classification by matching the expected pattern of SNPs in Walker *et al*. [12].

### Wide and Deep Neural Network Model

Wide and deep neural networks (WDNN) marry two successful models, logistic regression and deep multilayer perceptrons (MLP), to leverage the strengths of each approach. In WDNNs, a ‘wide’ logistic regression model is trained in tandem with a ‘deep’ MLP and the two models are merged in a final classification layer, allowing the network to learn useful rules directly from the input data and higher level nonlinear features. For genomic data, the logistic regression portion of network can be thought of as modeling the additive portion genotype-phenotype relationship, while the MLP models the nonlinear or epistatic portion. We implemented a multidrug wide and deep neural network [22] with three hidden layers each with 256 rectified linear units (ReLU) [35], dropout [36], batch normalization [37], and L2 regularization (Figure 1). L2 regularization was applied on the wide model (which is equivalent to the well-known Ridge Regression model) [38], the hidden layers of the deep model, and the output sigmoid layer. The network was trained via stochastic gradient descent using the Adam optimizer.

**Fig 1.**
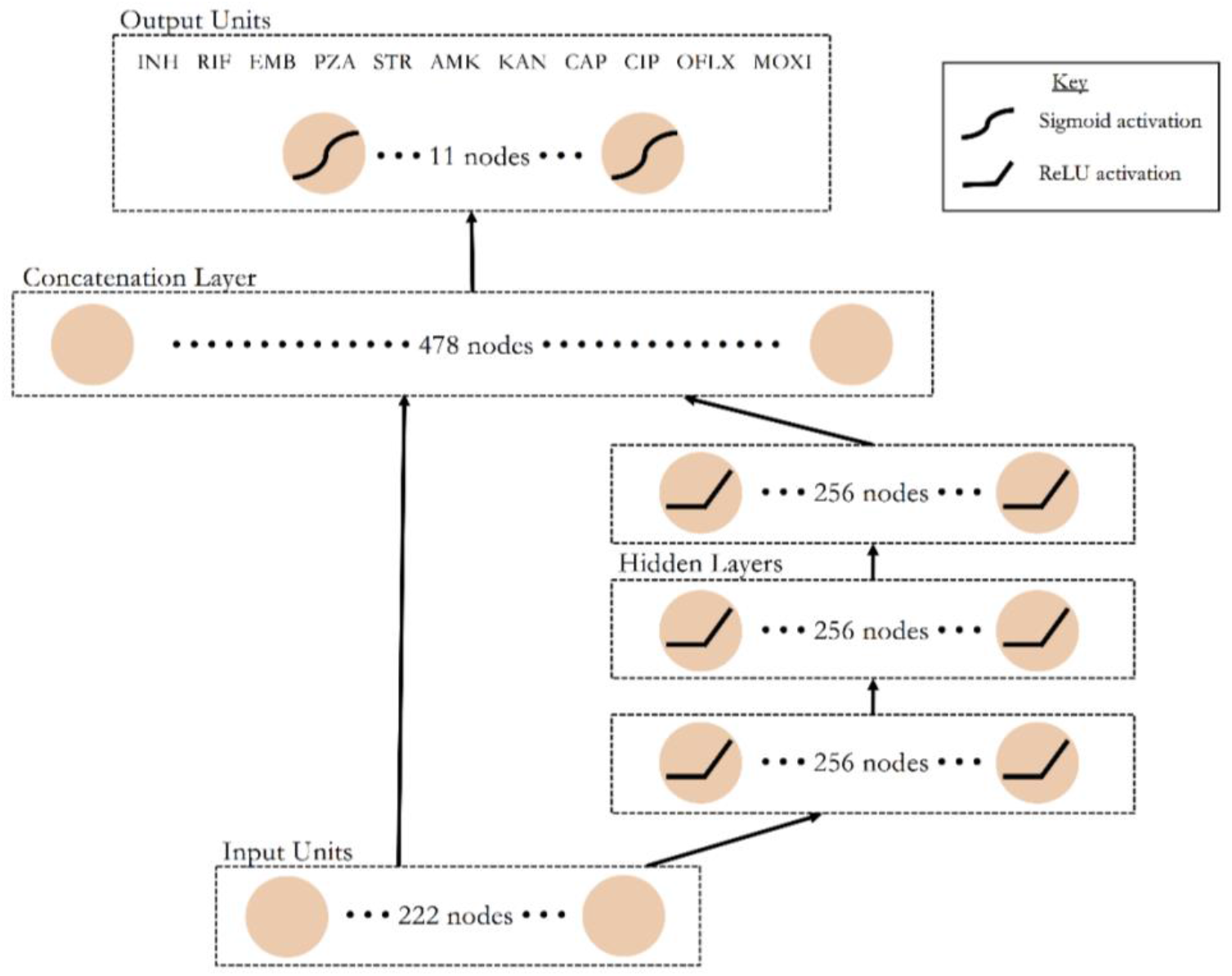
A schematic of the multidrug wide and deep neural network architecture. *Data flows from bottom to top through the wide (left) and deep (right) paths of the neural network. Nonlinear transformations, where applied, are depicted on the corresponding nodes. Each of the 11 nodes in the output layer represents resistance status predictions in all MTB isolates for one of the 11 anti-tuberculosis drugs*.

The MD-WDNN was trained simultaneously on resistance status for all 11 drugs, including ciprofloxacin. Each of the 11 nodes in the final layer represented one drug and its output value was the probability that the MTB isolate was resistant to the corresponding drug. We constructed single-drug WDNNs (kSD-WDNN and SD-WDNN) with the same architecture as the multidrug model except for the structure of the output layer, which predicts for one drug.

The MD-WDNN utilized a loss function that is a variant of traditional binary cross entropy. Our dataset had a missing resistance status for some drugs in the MTB isolates, so we implemented a loss function that did not penalize the model for its prediction on drug-isolate pairs for which we did not have phenotypic data. Due to imbalance between the susceptible and resistant classes within each drug, we adjusted our loss function to upweight the sparser class according to the susceptible-resistant ratio within each drug. Thus, the final loss function was a class-weight binary cross entropy that masked outputs where the resistance status was missing.

### Baseline Models

In addition to the multidrug and single drug wide and deep neural networks, we implemented three other classification models – a single drug random forest, a single drug L2 regularized logistic regression, and a deep MLP with the same architecture as MD-WDNN without the ‘wide’ logistic regression model. The deep MLP was used as a baseline to justify the importance of the regression portion of the WDNN architecture.

### Training and Model Evaluation

The MD-WDNN, SD-WDNN, random forest, regularized logistic regression, and deep MLP were trained on predictors in the dataset present in at least 30 MTB isolates. The kSD-WDNN trained on mutations based on preselected genes, as described above. The MD-WDNN (Common Mutations) trained only on mutation predictors without the derived features to measure the effect on accuracy of these derived features.

We repeated ten-fold cross validation 5 times, for a total of 50 different validation sets, to train the models and evaluate performance. The SD-WDNN, kSD-WDNN, random forest, and regularized logistic regression models were stratified by class label to address imbalances between resistance and susceptible classes, as they were all single task classifiers. Final model (MD-WDNN only) performance was validated through an independent validation set.

We reported AUC for all the models during repeated 10-fold cross-validation. We ranked the models from 1-7 (best to worst) based on AUC for each of the 10 anti-tuberculosis drugs and reported an average rank for each model. For the MD-WDNN, we reported two pairs of sensitivity and specificity performance on the independent validation set. We reported the first pair by determining a probability threshold to maximize the sum of specificity and sensitivity for each drug. For the second pair, we determined the probability threshold to maximize sensitivity given that the specificity is at least 90%. The 90% specificity threshold stems from the value assessment that over-diagnosis of antibiotic resistance is more harmful than under-diagnosis due the treatment toxicity and side effects, *e.g*. renal failure and hearing loss, for the drugs used in antibiotic resistant cases. During repeated ten-fold cross-validation, we reported the mean and confidence of specificity and sensitivity based on validation set results across the 50 different validation sets.

### MTB isolate visualization using t-SNE

We applied *t*-distributed Stochastic Neighbor Embedding (*t*-SNE), a method for visualizing data with high dimensionality [39], to two datasets: (1) the set of input predictors (common mutations and derived categories) and (2) the final output layer of the MD-WDNN. The genetic markers, originally in 222 dimensions, and the final layer weights, originally in 11 dimensions, were each extracted from the MD-WDNN and projected onto two dimensions. Each point represented one MTB isolate and was colored based on its phenotypic status for each drug. The lineage clustering was also overlaid on the *t*-SNE plots to determine the effect of lineage on the two different representations of isolates.

### Importance of MTB genetic variants to drug resistance

We examined predictor importance to resistance by analyzing the prediction outputs of the MD-WDNN and the presence or absence of mutations through a permutation test. We permuted the resistance labels and calculated the distribution of following difference:

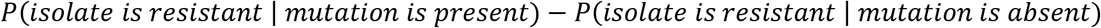

where P(isolate is resistant | mutation is present) is the MD-WDNN’s probability of resistance for a given mutation. We then compared the actual differences with the permuted differences. The sampling distribution included 100,000 randomized permutations per mutation and the actual differences were evaluated at a significance level of *α* = 0.05 corrected for multiple comparisons. We conducted the permutation test for each predictor (common mutations or derived categories) that was present in at least 30 MTB isolates. We focused on the common mutations and derived categories that were correlated with resistance to anti-tuberculosis drugs.

### Implementation Details

All WDNN and MLP model implementations used the Keras 1.2.0 library in Python 2.7 with a TensorFlow 0.10.0 backend. The random forest and regularized logistic regression classifiers were implemented with Python Scikit-Learn 0.18.1. The isolate diversity analysis was implemented using R 3.4.0, the *t*-SNE analysis used the Rtsne 0.13 package in R, and the permutation tests were implemented in Python 2.7. All models were trained on a NVIDIA GeForce GTX Titan X graphics processing unit (GPU). Hyperparameters are available in Supplementary Table S7. All analysis code and input data files are openly available at https://github.com/farhat-lab/wdnn.

## Results

### Data Processing

The pooled data from the WHO network of supranational reference laboratories and the ReSeqTB knowledgebase [8,23] used in training the initial model included 3,601 MTB isolates. All of the anti-tuberculosis drugs had a higher proportion of susceptible isolates compared to resistant isolates, ranging from 53.0% to 88.1% susceptible for the different drugs. Ofloxacin was tested on the smallest number of isolates at a total of 739. All other drugs were tested in at least 1,204 isolates, with rifampicin tested in 3,542 isolates and isoniazid in 3,564 isolates (Supplementary Table S1).

The independent validation set contained 792 MTB isolates, with 198 to 736 of these isolates tested for each of the 10 drugs (Supplementary Table S2). Because ciprofloxacin had limited phenotypic availability in the independent validation set and predictive performance could not be validated, we did not include performance for ciprofloxacin resistance.

We found 6,342 different insertions, deletions, and single nucleotide polymorphisms (SNPs) in 30 promoter, intergenic, and coding regions of the MTB isolates’ genomes. Of these variants, 166 were present in at least 30 of the 3,601 isolates and were used as predictors in Dataset 1. Of the 3,445 variants found in fewer than 30 isolates, we aggregated the variants into 141 derived categories (see Methods) and used 56 of them, those present in at least 30 isolates, as predictors. The final model used 222 total predictors in training and subsequent analyses.

### Evaluation of MTB isolate diversity

Sequence data from 33 genetic lineage markers (Supplementary Table S3) were available in all 3,601 isolates and were used to measure genetic distance between isolates [12]. The isolates fell into five well-defined clusters that corresponded to MTB’s known genetic lineages. All 5 lineages were well represented with 632 from the Euro-American Latin America Mediterranean sub-lineage, 1,501 from other Euro-American sub-lineages, 331 from the Indo-Oceanic or *Mycobacterium africanum*, 643 from the Central Asian lineage, and 494 from the East Asian lineage (Figure 2). The input layer *t*-SNE coordinates also largely recapitulated the genetic clustering due to lineage (Supplementary Figure S1), illustrating that the largest genetic differences between isolates were related to lineage. On the other hand, overlying *t*-SNE coordinates for the MD-WDNN’s probabilistic representation (Supplementary Figure S2) confirmed that the MD-WDNN’s prediction of phenotype was not biased by lineage related variation.

**Fig 2.**
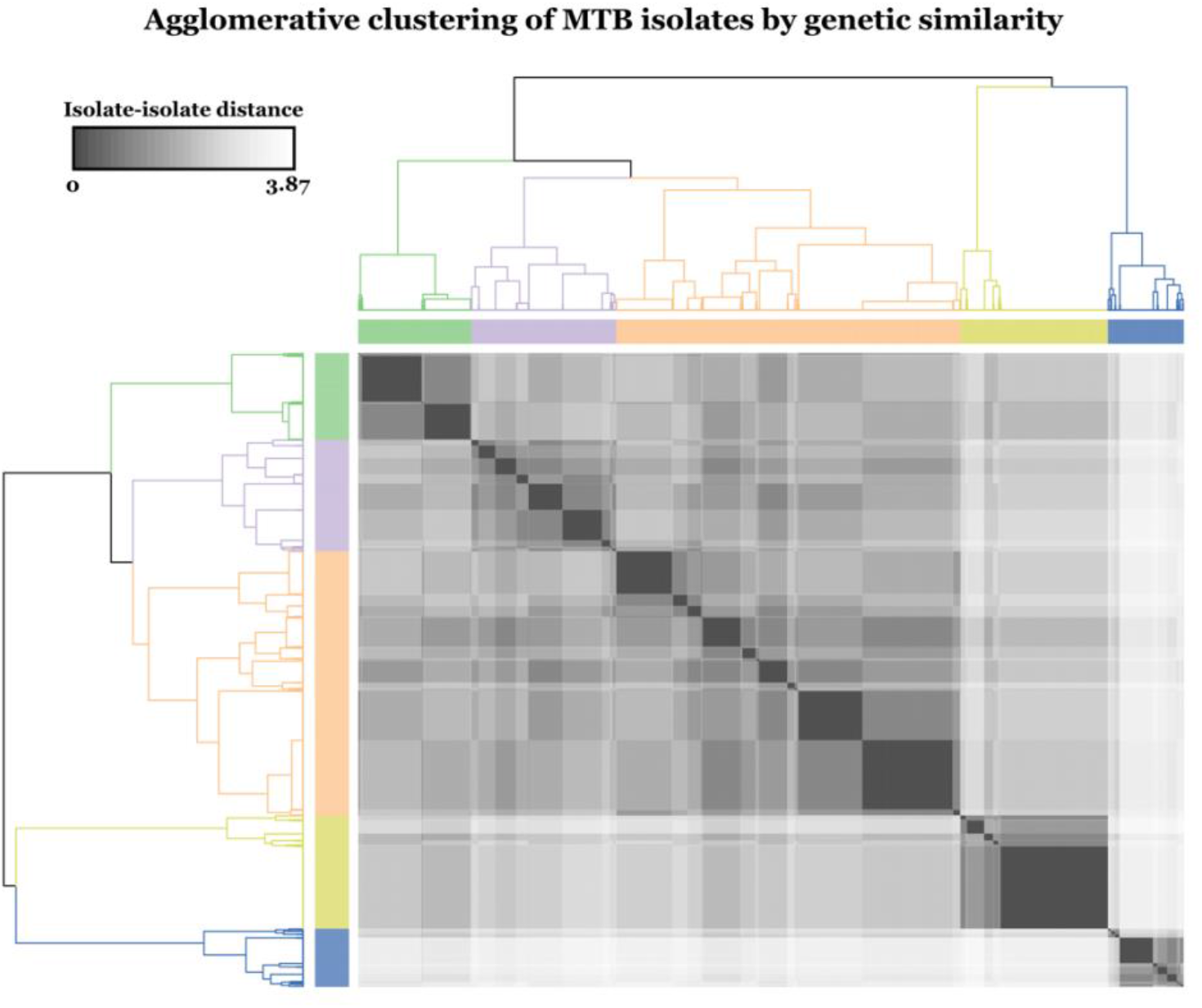
Agglomerative clustering of MTB isolates by genetic similarity. *We used known lineage-defining mutations to calculate isolate-isolate Euclidean distances, which is shown in the heat map. Using these distances of the lineage-defining mutation vectors between isolates, we applied Ward’s method of hierarchical clustering to construct the dendrogram and determine the five lineage clusters. The dendrogram is colored by the corresponding isolate genetic lineage. Green: East Asian, Purple: Euro-American (Latin America Mediterranean-LAM sublineage), Orange: Euro-American (other sublineages than LAM), Yellow: Central Asian, Blue: Indo-oceanic & M. africanum*.

### Comparison of model predictive performance

Although random forest, logistic regression and simpler versions of the deep learning models performed well (AUCs by repeated ten-fold cross validation in Supplementary Table S4), the MD-WDNN trained on the largest number of genetic predictors ranked highest in performance across both first and second line drugs (average rank of 2.1). The next best ranked models were the deep MLP, regularized logistic regression, and random forest respectively, with average ranks of 3, 3.4, and 3.8, across all ten drugs. The largest step up in AUC was observed between the kSD-WDNN, the model trained using genetic regions known to be causative of resistance, and any of the models trained on the full predictor set (Figure 3, Supplementary Table S4). For the second line drugs, the average AUC was 0.879 for kSD-WDNN vs 0.936 for the MD-WDNN. The importance of using rare genetic variation in predicting resistance is highlighted by the loss of performance seen with the WDNN built without the derived variables (Figure 3). This was most notable for the drug pyrazinamide.

**Fig 3.**
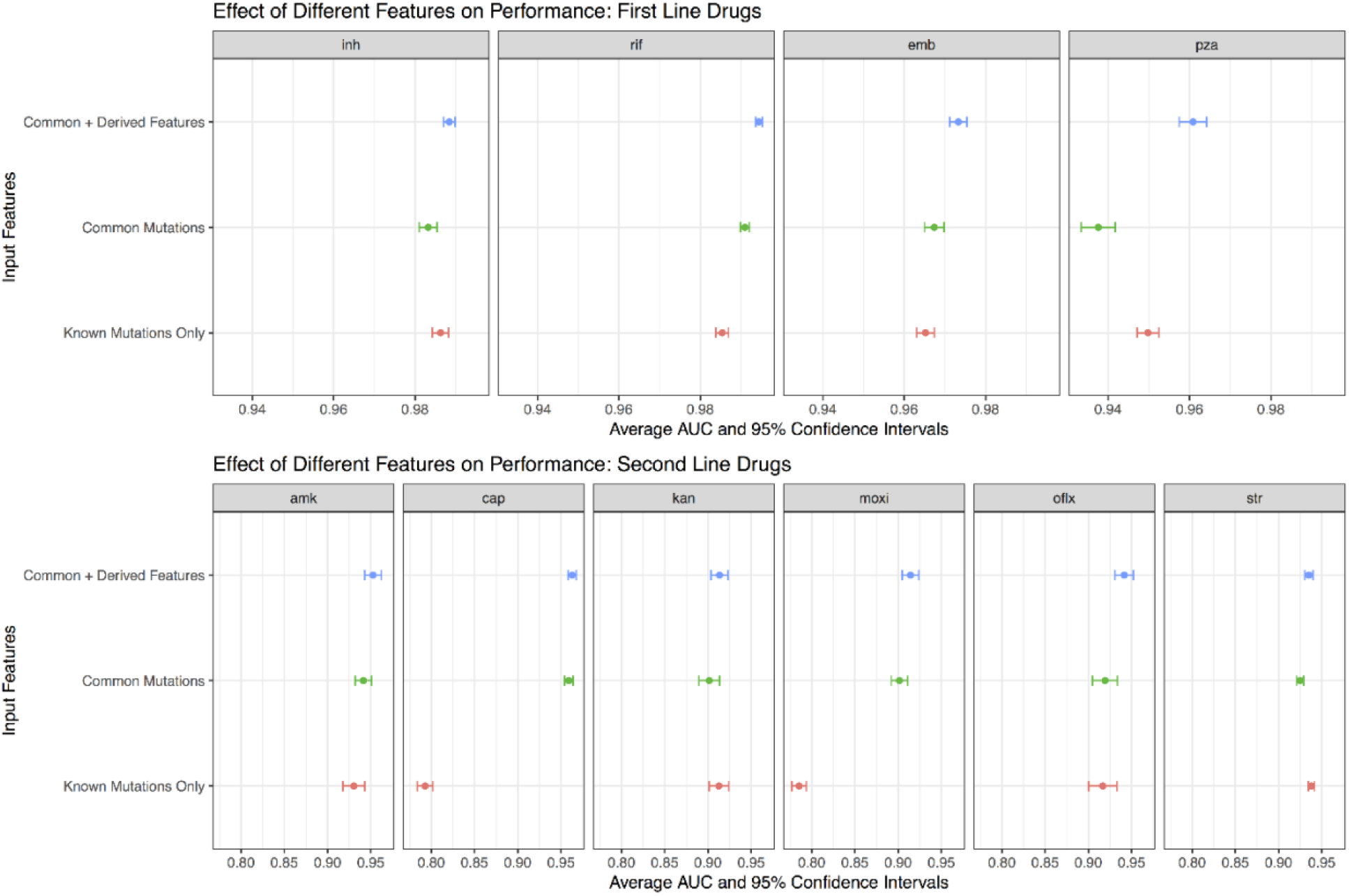
Comparison of tuberculosis drug resistance predictive performance based on input feature set. *Area under the ROC curve classification performance during repeated cross-validation for the MD-WDNN trained on all features (mutations and derived categories), MD-WDNN trained on just mutations (MD-WDNN (Common Mutations)), and the single drug WDNN trained on mutations and derived categories occurring in genes known to be resistant determinants for each drug (kSD-WDNN)*.

Next, we compared the performance of a single drug model to the multidrug version. The predictive performance of the MD-WDNN and the SD-WDNN during repeated cross-validation are shown in Figure 4. The average AUC for the SD-WDNN was 0.978 for first-line drugs and 0.928 for second-line drugs; the multidrug architecture of the MD-WDNN resulted in a higher average AUC for both first-line and second-line drugs. For most drugs, we see modest improvements in performance for the MD-WDNN compared with the SD-WDNN, with larger gains observed for the drugs kanamycin and ofloxacin.

**Fig 4.**
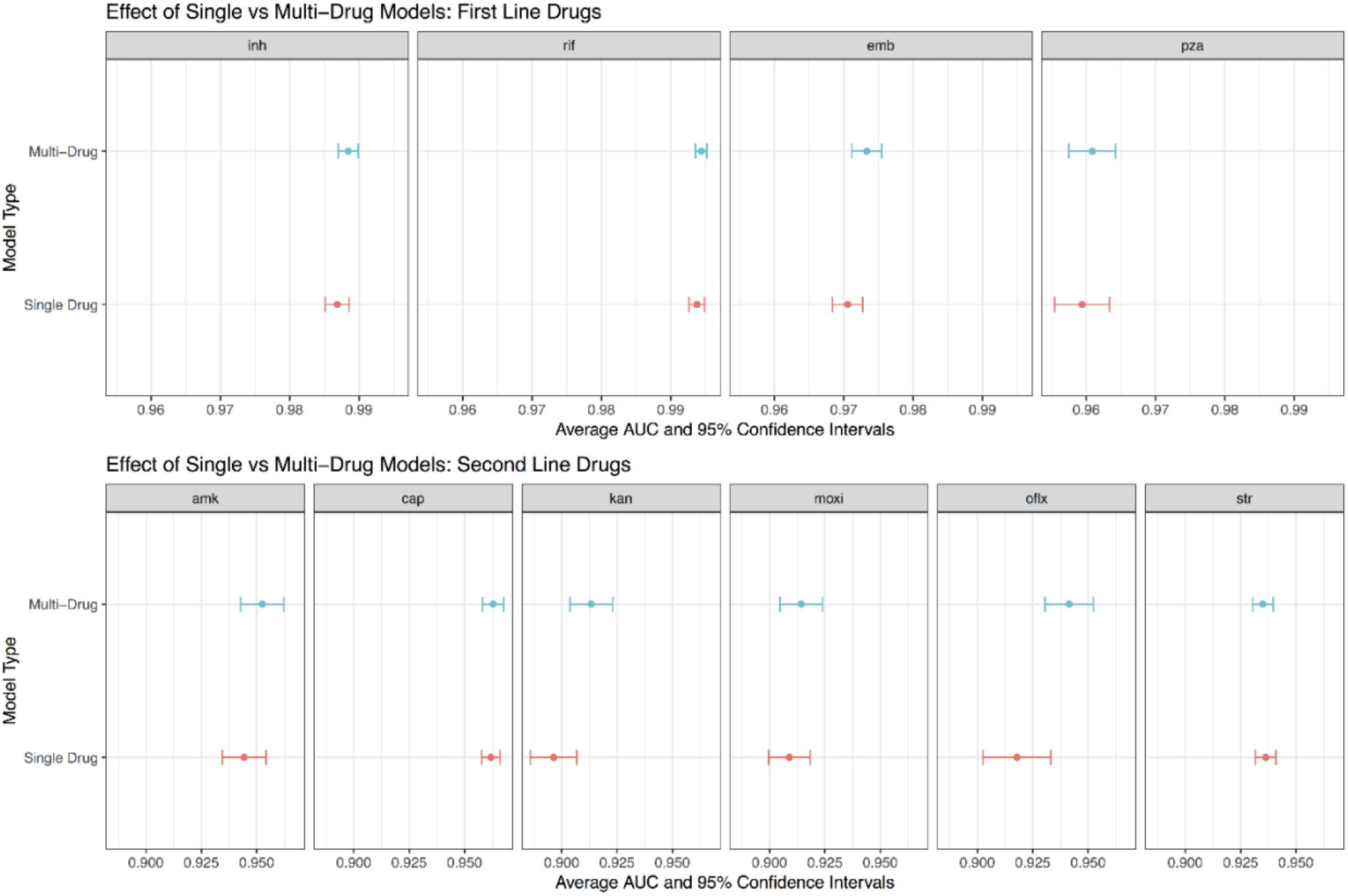
Comparison of tuberculosis drug resistance predictive performance between single drug and multidrug models. *Area under the ROC curve classification performance during repeated cross-validation for the multidrug WDNN predicting resistance for all drugs simultaneously and for the single drug WDNN*.

The cross-validation results above supported the choice of the MD-WDNN as the final model and we proceeded to validate its performance on an independent data set. The ROC curves for the MD-WDNN on the independent validation data across the 10 anti-tuberculosis drugs are shown in Figure 5, illustrating the different sensitivity and specificity performance values at probability thresholds between 0 and 1. Table 1 shows the AUC corresponding to the ROC curves for each drug. The average AUCs were 0.937 for first-line drugs and 0.891 for second-line drugs on an independent validation set, which were slightly lower than the AUCs during repeated cross-validation (AUC = 0.979 for first-line drugs, AUC = 0.936 for second-line drugs). Sensitivities, specificities, and the corresponding probability threshold, which was chosen to maximize the sum of sensitivity and specificity, are also shown in Table 1. The average sensitivities and sensitivities, respectively, on the independent validation set were 87.9% and 92.7% for first-line drugs and 82.0% and 90.1% for second-line drugs. Sensitivity and specificity values for the second probability threshold, which maximizes sensitivity given that specificity is at least 90%, are available in Supplementary Table S5.

**Fig 5.**
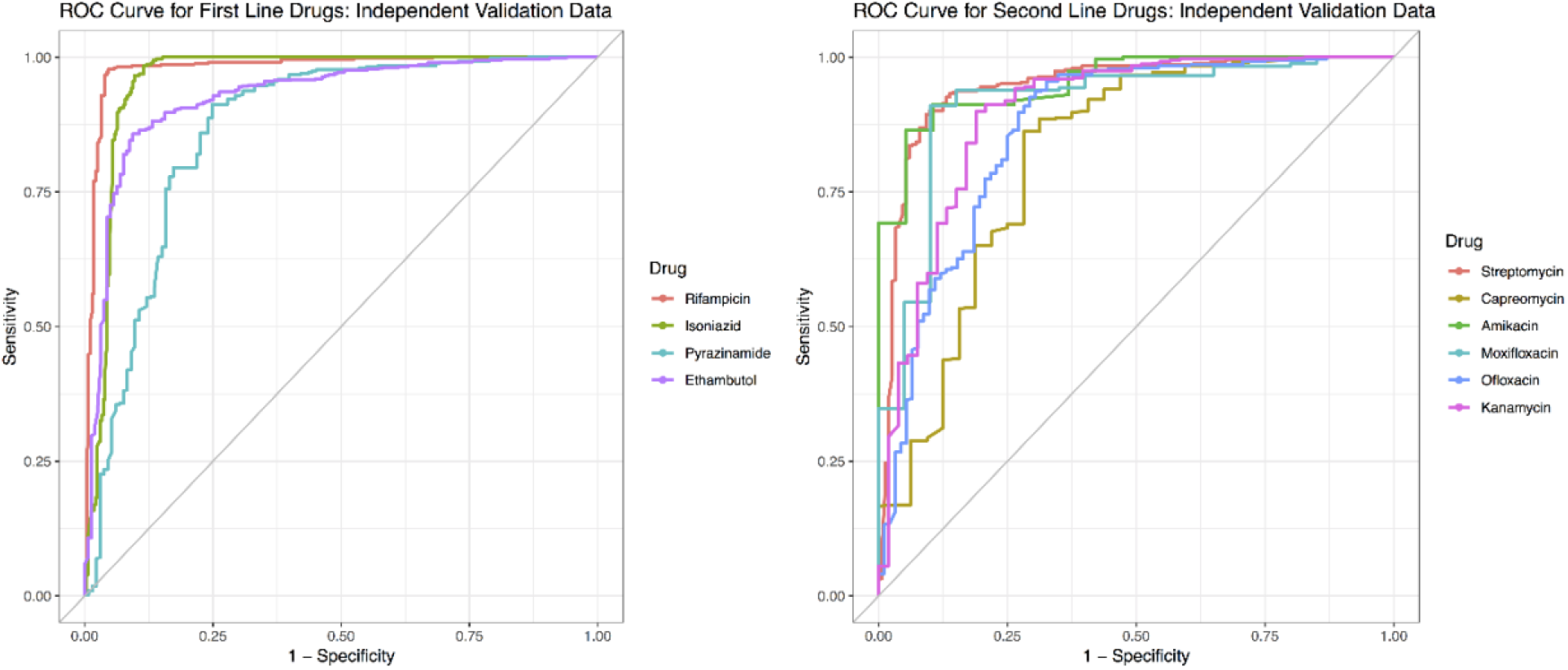
Tuberculosis drug resistance ROC performance curve of the MD-WDNN. *A ROC plot of MD-WDNN predictive performance on the independent validation set for first-line (left) and second-line (right) anti-tuberculosis drugs*.

**Table 1.** Tuberculosis drug resistance predictive performance of the MD-WDNN. *Area under the ROC curve classification performance on the independent validation set. We also report sensitivity and specificity performance with the probability threshold chosen to maximize the sum of sensitivity and specificity for all anti-tuberculosis drugs*.

| Drug | AUC | Sensitivity | Specificity | Threshold |
| --- | --- | --- | --- | --- |
| Rifampicin | 0.982 | 95.4% | 97.8% | 0.35 |
| Isoniazid | 0.959 | 90.3% | 96.4% | 0.09 |
| Pyrazinamide | 0.883 | 75.2% | 91.2% | 0.32 |
| Ethambutol | 0.922 | 90.6% | 85.6% | 0.40 |
| Streptomycin | 0.942 | 90.1% | 89.6% | 0.26 |
| Capreomycin | 0.808 | 71.9% | 85.7% | 0.23 |
| Amikacin | 0.950 | 89.5% | 90.8% | 0.20 |
| Moxifloxacin | 0.902 | 90.0% | 91.0% | 0.39 |
| Ofloxacin | 0.866 | 69.6% | 93.7% | 0.57 |
| Kanamycin | 0.879 | 81.1% | 89.6% | 0.32 |

### MTB isolate visualization using t-SNE

A popular way to visualize the components of a deep learning model is the *t*-distribution stochastic neighborhood embedding (*t*-SNE) method, which is a nonlinear dimensionality reduction technique [39]. We applied *t*-SNE separately to (1) the input genetic predictors and (2) the MD-WDNN predictions. *t*-SNE on the input genetic markers showed well-defined clusters, and each cluster contained both susceptible and MDR isolates with little discernable pattern of resistance classification (Supplementary Figure S3). Contrarily, Figure 6 demonstrates clear separation by the MD-WDNN’s output representation between resistant and susceptible isolates, consistent with our measurements of high model sensitivity and specificity. The *t*-SNE plots also demonstrates the multitask WDNN’s ability to classify resistance across multiple drugs, separating them into nested groups of pan-susceptible isolates, followed by mono-INH resistant isolates, multidrug resistant isolates, pre-XDR isolates, and XDR isolates, which is consistent with the order of administration of the drugs clinically as well as the usual order of MTB drug resistance acquisition [40]. The second-line injectable drugs, AMI, CAP, and KAN, also show similarly-classified clusters, highlighting the well-known moderate level of cross resistance between them. We also observe this among the fluoroquinolones despite the fact that fewer isolates were tested for resistance to these agents [41].

**Fig 6.**
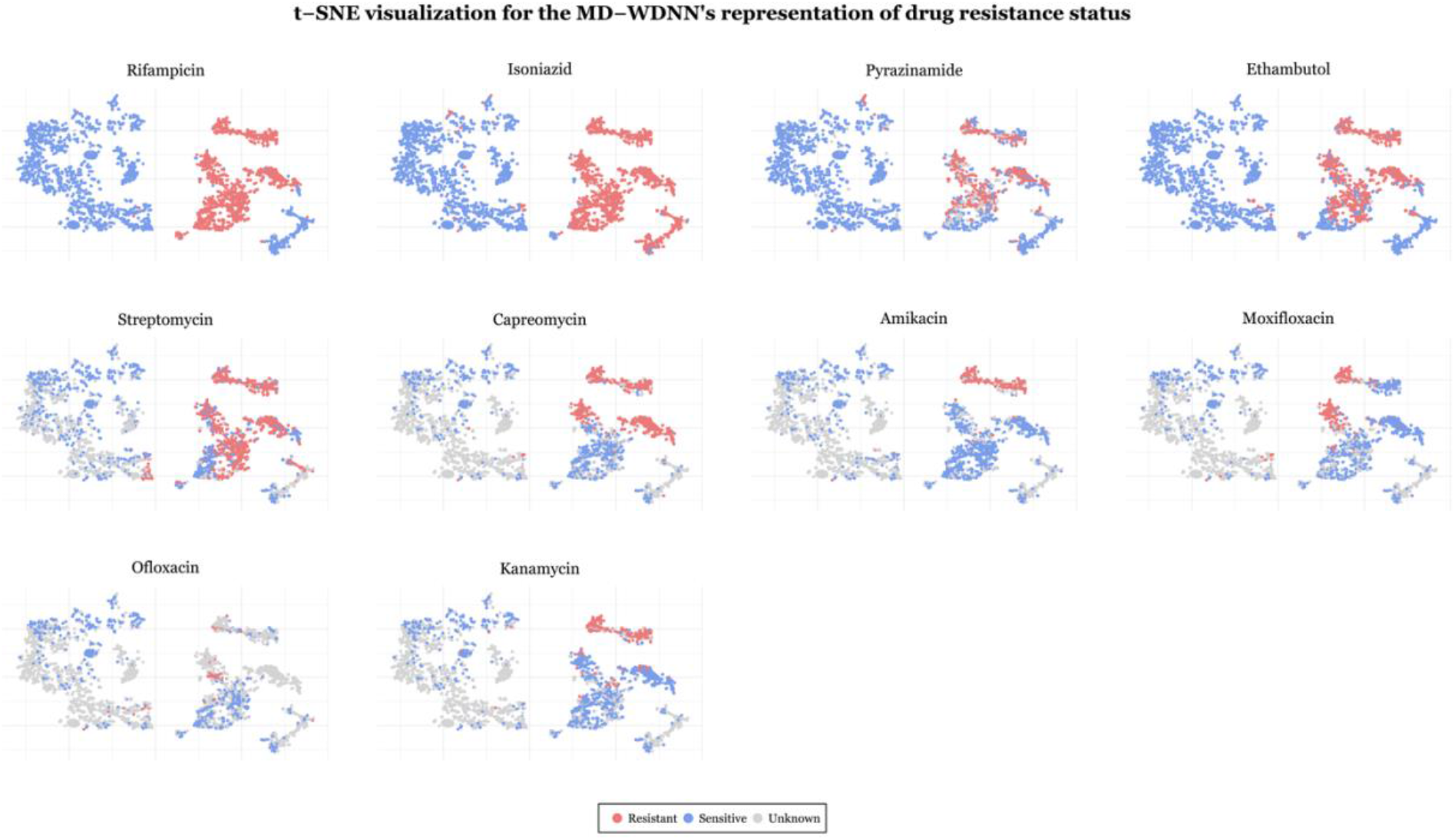
t-SNE visualization for the final output layer of the MD-WDNN. *The final layer predictions, originally in 11 dimensions, were projected onto two dimensions. Each point is an MTB isolate, colored according to its resistance status with respect to the corresponding drug*.

### Importance of MTB genetic variants to drug resistance

All 222 predictors were tested for importance to resistance to each of the 10 drugs through a permutation test as described in the methods section. The first-line anti-tuberculosis drugs had the largest numbers of significant ‘resistance predictors’: rifampicin (143 predictors), isoniazid (144 predictors), pyrazinamide (132 predictors), ethambutol (140 predictors), as well as the second-line drug streptomycin (140 predictors).

Figure 7 illustrates the number of significant predictors per drug and the predictor intersections among different drug subsets. There were 37 drug subsets that shared at least one resistance predictor. The largest subset was of 10 anti-tuberculosis drugs that shared 69 resistance predictors. Subsets of drugs that included a second line injectable drug and shared at least two predictors consistently included both INH and RIF. This is consistent with previous findings that MTB isolates acquire resistance to first-line drugs before second-line drugs [40] and indicates that the multidrug model was able to capture these relationships. The subset of fluoroquinolones shared 3 resistance-correlated predictors not found in other first-line or second-line drugs, which is expected given that fluoroquinolones have a mechanism of action that differs from those of first-line and second-line drugs [42].

**Fig 7.**
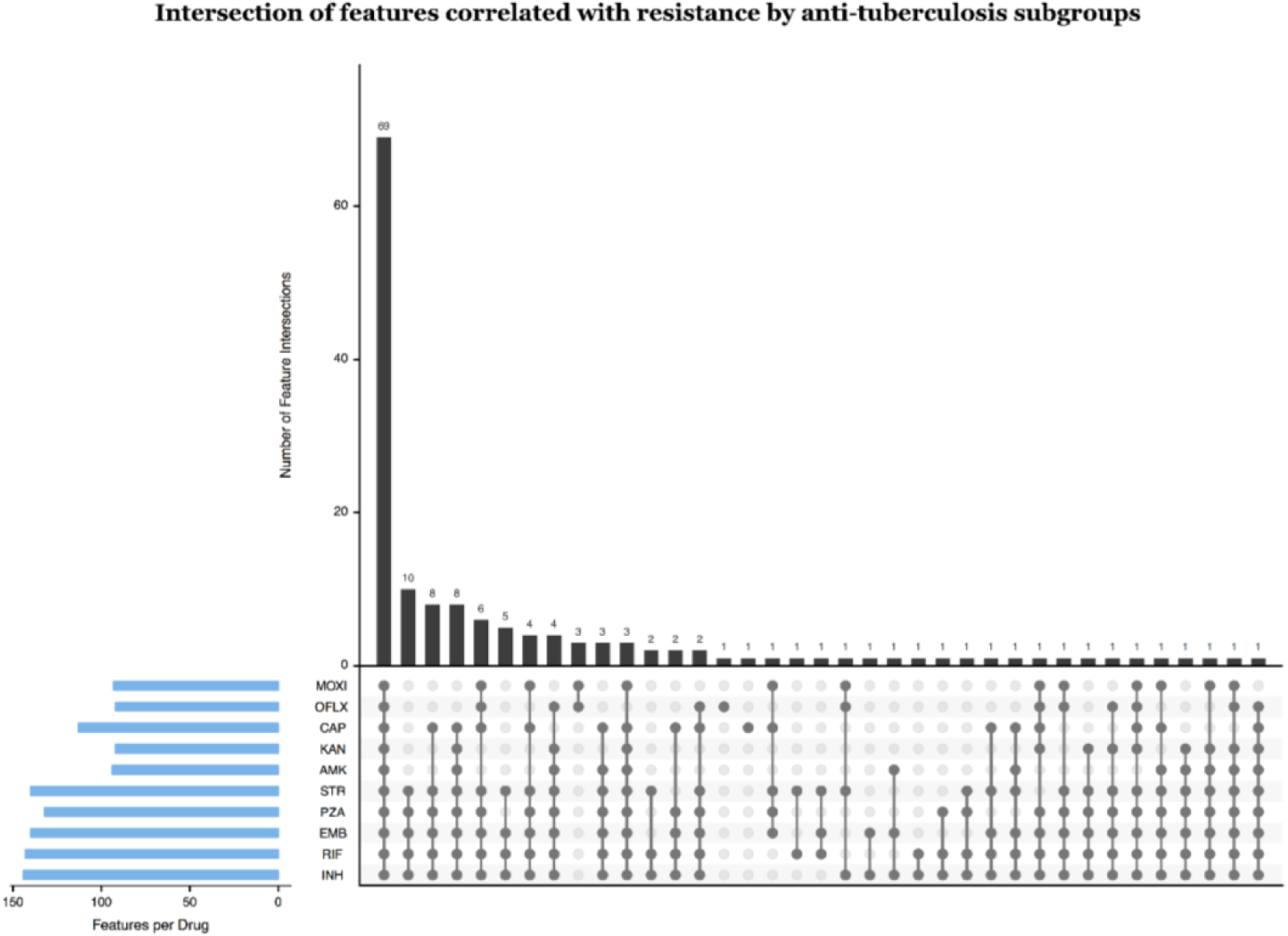
Intersection of predictors correlated with resistance by anti-tuberculosis drug subgroups. *We permuted the resistance labels and calculated the distribution of the difference, P(isolate is resistant* | *mutation is present) – P(isolate is resistant* | *mutation is absent). We show the number of mutations per subgroup of drugs ordered from most to least mutations per subgroup. Number of significant predictors per drug is also shown*.

## Discussion

The primary aim of this study was to construct a highly accurate model of drug resistance using genomic data. A few prior studies have utilized algorithmic or machine learning methods using MTB genomic data to account for the complex relationship between genotype and drug resistance [8,12,13,27]. We demonstrate here that the multidrug WDNN approach outperforms our previously reported random forest model [8]. Compared to one study that used a direct association (DA) algorithm, the MD-WDNN presented here offers improvement in sensitivity and specificity for the majority of drugs when prediction is attempted on all isolates, including those with rarer and not previously observed variants [12]. One study used single-task machine learning, demonstrating the validity of this approach for identifying MDR and XDR-TB, but was limited by the use of a dataset with a low number of MDR isolates (n=81) and even lower numbers of isolates resistant to drugs other than RIF and INH (n ranging from 19 to 59), raising concerns about generalizability [13].

Our approach has several features which are important to its success. First, we included all variants in the set of 30 genetic loci as potential predictors of resistance to any drug and did not subset the variants according to *a priori* knowledge of causative relationships between genetic loci and drugs. The performance gains offered by this more ‘permissive’ approach were considerable especially for the second line drugs, and the first-line ‘sterilizing’ drug pyrazinamide [43]. Second, the multidrug structure allows drugs which have less phenotypic data to borrow information about resistance pathways from drugs that have higher numbers of phenotyped isolates. Additionally, the wide and deep structure allows us to include prior information about the genetic etiology of MDR and XDR, as it is known that both individual markers and gene-gene interactions confer resistance [9–11]. The wide portion of the network allows the effect of individual mutations (*e.g*. marginal effects) to be easily learned, while the deep portion of the network allows for arbitrarily complex epistatic effects to influence the predictions. The model presented here is the first multitask tool that predicts resistance for 10 anti-tuberculosis drugs simultaneously with state-of-the-art performance.

Multitask architectures in deep learning have not been used widely in pharmaceutical and drug-related industries due to many barriers, including the difficulty of implementing a high-quality deep multitask network [44]. However, past multitask deep learning algorithms have seen success over traditional single task baseline models, such as in applications to drug discovery and studying gene regulatory networks [44–46]. In addition, multitask neural networks have been shown to have larger performance gains over single task models when using smaller datasets [47,48]. Our study demonstrates the utility of the multidrug framework coupled with more permissive inclusion of genetic variables for predicting antibiotic resistance. Although the gains were modest for several drugs, in the case of kanamycin and ofloxacin the performance gains were larger. This is an important advance as second line injectables and fluoroquinolones are cornerstone agents for the treatment of MDR-TB treatment, and accurate prediction of susceptibility to these agents is key in determining a patient’s candidacy for the recently recommended shortened MDR-TB regimen [49]. Prediction of resistance to second-line injectables has thus far been challenged by a limited genetic knowledge base and consequently limited sensitivity when using simple direct association approaches [12]. Thus, the use of a more complex model, such as our multidrug WDNN, appears to be justified at least for a subset of the anti-tuberculosis drugs.

The increase in predictive performance of the MD-WDNN over other models assessed may arise from a number of possibilities. First, phenotypic resistance data that was highly available in our dataset for certain drugs (*i.e*. RIF, INH, and EMB) served as a direct indicator for resistance to second-line injectables and fluoroquinolones. This is unlikely to be the only reason, as our *t*-SNE analysis shows clustering patterns specific to second-line injectable drugs and fluoroquinolones, and the validated model specificity for these drugs was robust. Second, mutations that do not necessarily confer resistance to particular drugs may be indicative of other genomic predictors, thereby serving as a reliable predictor for resistance. Because of the large intersection of mutations (Figure 7) for all anti-tuberculosis drugs, again this cannot be the sole explanation for the performance differences. The correlative effect of mutations can be treated as a positive feature in the multidrug architecture. On the other hand, the potential lack of causation also requires care when using the predictive model. Third, there may exist mutations, especially those that are less frequent, that are not yet known to confer resistance to particular anti-tuberculosis drugs but were captured by the multidrug WDNN thereby improving performance.

Understanding the improved performance of our wide and deep neural network is a particularly difficult task due to the architectural complexity and lack of visualization tools in deep learning [50,51]. However, we gained some insights by comparing the *t*-SNE visualizations of the input data and model predictions. This confirmed that the model is not biased by the isolate genetic lineage structure, a recognized confounder of genotype and phenotype relationships in MTB, even though antibiotic resistance was unevenly distributed among the different lineages. The fact that the MD-WDNN was not biased by isolate lineage structure is also demonstrated by the high performance of the model on the independent data set.

The translation of our deep learning approach is also a function of advancements in whole genome sequencing and accessibility to more MTB isolate data. Improvements in whole-genome sequencing technologies have significantly reduced costs [52], allowing for more routine whole genome sequencing in MTB isolates [53]. The prediction time for MTB drug resistance depends primarily on the sequencing turnaround time, which is significantly shorter than phenotypic susceptibility testing [54]. In addition, as more routine sequencing increases the amount of MTB isolate data, our deep learning model can be rapidly updated as the datasets become accessible. We expect that as more data are incorporated, the sensitivity and specificity gap in second-line injectable drugs and fluoroquinolones will become smaller.

We acknowledge some limitations of our study. First, one source of bias could be errors during phenotyping, as susceptibility testing for some drugs has been shown to have low reproducibility and high variance [55]. However, we used strains with phenotypic data measured at national or supranational TB reference laboratories following strict quality control or carefully curated from research and reference laboratories [8,23]. Beyond technical or laboratory limitations in testing, certain resistance mutations, especially for ethambutol and second-line drugs, may result in minimum inhibitory concentrations (MICs) very close to the clinical testing concentration, which may result in lower sensitivity and specificity [56] when predicting a binary resistance phenotype. The use of MIC data for building future learning models may help circumvent this. Second, we only included mutations that occurred in >0.8% (30 of 3,601 isolates) individually or when aggregated with other rare variants in the same gene or intergenic region. Although we may have missed some important predictors, this threshold amounted to only ignoring variants that are very rare in a diverse sample of MTB genomes with good representation from the 4 major genetic lineages. Third, we did not include third-line antituberculosis drugs such as cycloserine or para-aminosalicylic acid due to the lack of phenotypic data.

In summary, we present a new deep learning architecture to identify the resistance of MTB isolates to 10 anti-tuberculosis drugs from whole genome sequencing data. The wide and deep neural network achieved state-of-the-art performance on a large, aggregated TB dataset, demonstrating the efficacy of deep learning as a diagnostic tool for MTB drug resistance. The WDNN represented the first multidrug model to our knowledge that incorporated a high number of genotypic predictors known to be important to determining resistance for one or more included drugs. Further work identifying the key processes of deep learning will not only allow for improved predictive performance but may also give us a greater understanding of the biological mechanisms underlying drug resistance in MTB isolates.

